# Measuring Perceptual Distance of Organismal Color Pattern using the Features of Deep Neural Networks

**DOI:** 10.1101/736306

**Authors:** Drew C. Wham, Briana Ezray, Heather M. Hines

## Abstract

A wide range of research relies upon the accurate and repeatable measurement of the degree to which organisms resemble one another. Here, we present an unsupervised workflow for analyzing the relationships between organismal color patterns. This workflow utilizes several recent advancements in deep learning based computer vision techniques to calculate perceptual distance. We validate this approach using previously published datasets surrounding diverse applications of color pattern analysis including mimicry, population differentiation, heritability, and development. We demonstrate that our approach is able to reproduce the biologically relevant color pattern relationships originally reported in these studies. Importantly, these results are achieved without any task-specific training. In many cases, we were able to reproduce findings directly from original photographs or plates with minimum standardization, avoiding the need for intermediate representations such as a cartoonized images or trait matrices. We then present two artificial datasets designed to highlight how this approach handles aspects of color patterns, such as changes in pattern location and the perception of color contrast. These results suggest that this approach will generalize well to support the study of a wide range of biological processes in a diverse set of taxa while also accommodating a variety of data formats, preprocessing techniques, and study designs.

## INTRODUCTION

Organisms display a rich variety of patterns and colors in nature partly because selection operates on perceptual form, while drift prevents its perfect cohesion over time and space (Hoffman et al. 2006, Gray and McKinnon 2007). The opportunities for selection to act on organismal color patterns are diverse, including camouflage (Ruxton et al. 2004, Merilaita 2003, Cott 1940), warning signals (Chouteau et al. 2016, Ruxton et al. 2004, Sherratt 2002, Mallet and Singer 1987), thermoregulation (Brakefield 1984, Stiles 1979, Williams 2007), mate recognition (Couldridge and Alexander 2002, Merrill et al. 2012), sexual selection (Endler 1983, Endler and Basolo 1998, Jiggins et al. 2001), and attraction of symbiotic partners (e.g. relationship between flowers and pollinators) (Schiestl and Johnson 2013). Such selection has driven many systems to attain high levels of color pattern variation, with exemplary systems including invertebrates such as spiders (Cotoras et al. 2016), *Heliconius* butterflies (Turner 1981, Sheppard et al. 1985, Mallet and Joron 1999), velvet ants (Wilson et al. 2012, Wilson et al. 2015), and bumble bees (Williams 2007, Ezray et al. 2019); vertebrates such as hamlet fish (Thresher 1978), poison frogs (Symula et al. 2001), coral snakes (Cox and Rabosky 2013), birds (Endler and Thery 1996, Siefferman and Hill 2005) and primates (Bradley and Mundy 2008); as well as flowering plants (Rauscher 2008), like *Mimulus* (Cooley and Willis 2009), *Penstemon* (Wilson et al 2004) and morning glories (Baucom et al 2011).

The study and quantification of variations in color form in nature has been, and continues to be, fundamental to studies of evolution, ecology, and genetics. The quantification of the relationship between color patterns, however, is a uniquely difficult problem because color patterns can exhibit many axes of variation (Endler and Mappes 2017, Van Belleghem et al. 2017, Weller and Westneat 2018, Maia et al. 2019). These include variation in color hue, saturation, and brightness, as well as variations in the distribution of colors resulting in differences in shape and contrast. For studies seeking to accurately quantify organismal color patterns, these diverse facets of color and pattern must be simultaneously measured, weighted, and combined so as to emulate receiver perception in a meaningful way.

Developments in quantification techniques to measure individual color or pattern elements, as well as analytical processes combining multiple independently derived metrics into joint metrics, have provided researchers the ability to move beyond subjective descriptions of similarity. These methods have enabled a better understanding of ecological and evolutionary dynamics of color pattern formation in a variety of contexts. Some of these commonly used methods include pixel-by-pixel quantification (Williams 2007, Dittrigh et al. 1993), boundary strength analyses (Endler et al. 2017), adjacency analyses (Endler 2012), and color based analyses (Gawryszewski 2018, Weller and Westneat 2019). These methods quantify color patterns through a combination of salient spectral and spatial distribution properties (Endler 2012, Endler et al. 2017, Troscianko et al. 2017). Yet, several key limitations for these methods remain. Many of these methods rely on comparing traits by discrete body regions pixel-by-pixel. Such methods are sensitive to minor shifts in the location of color pattern elements, which may have little functional relevance to the receiver. Many methods convert color patterns into discrete presence-absence matrices, thus losing the continuous nature of pattern size, color, and spatial location. For instance, many of these methods homogenize color variation into small sets of discrete colors through K-means clustering, thus, ultimately losing the variable nature of color. Lastly, many of the applications developed either do not comprehensively address color and pattern simultaneously and/or they are not adaptable to other organisms (Weller and Westneat 2019, Van Belleghem et al. 2017). For this reason, several methods have been developed to integrate the various aspects of color pattern analysis into a single framework or analytical pipeline to measure each of these aspects either individually or collectively (Van Bellegham et al 2018, Maia et al. 2019, van den Berg et al. 2019).

To holistically quantify color pattern, an analysis needs to be able to concurrently assess color, pattern, and shape simultaneously. Transformative advancements in computer vision research, in particular the development and adoption of deep-learning based, convolutional neural networks (CNNs), have made this possible. Deep CNNs were designed to detect and categorize objects by learning patterns across a wide range of scales, typically starting at a 3×3 pixel frame and progressively expanding. The internal representation of an image in a CNN is calculated from every possible frame in an image for a series of frame sizes. These deep neural networks learn features (e.g., shapes, color patterns) across these frame sizes which discriminate between the groups or classes that they are trained to categorize. These CNNs have broad application because features learned during a sufficiently general classification task seem to be universally good features which adapt well to other classification tasks (Yosinski et al. 2014). Networks trained to categorize a large, diverse set of categories can often be adapted to classify new categories without having to learn new features. Among the most recent advancements in this research area is the discovery that distance measured in deep feature space correlates strongly with human judgement of perceptual distance (Johnson et al. 2016). Frequently called “Perceptual Loss”, this metric has underlied recent advancements in artistic style transfer (Johnson et al. 2016), image enhancement (Johnson et al. 2016, Ledig et al. 2017) and image synthesis (Wang et al. 2018, Karras et al. 2019). Recently, Zhang et al. (2018) subjected images to a variety of different types of distortions (e.g. warping, changes in saturation and hue, and blurring) and asked study participants to select the most similar images among sets of three. The results of this experiment confirmed that Perceptual Loss correlates strongly with human judgement, outperforming many other commonly used image similarity measures (Zhang et al. 2018). Zhang et al. (2018) further demonstrated that the correlation between Perceptual Loss and human judgment could be improved by re-weighting the deep features based on their correlation with human perceptual judgment in their experiment. This re-weighted Perceptual Loss metric was described as “Learned Perceptual Image Patch Similarity” (LPIPS) (Zhang et al. 2018).

Here, we provide a novel analytical pipeline that utilizes many of these recent advances in machine learning, including LPIPS, to enable quantification of similarity in color patterns. We validated the approach using several previously published datasets surrounding diverse applications of color pattern analysis including mimicry, population differentiation, heritability, and development. We demonstrate that when our approach is applied to these data it is able to recapitulate many of the previously reported findings. In contrast to the methods originally used to analyze these data, our approach does not require any task-specific training or input from the researcher. Thus, the approach is fully unsupervised. In many cases, we were able to reproduce the original findings directly from original photographs or plates with minimum standardization, avoiding the need for intermediate representation such as cartoonized images or trait matrices. Finally, we present two artificial datasets designed to highlight two challenging aspects of color pattern analysis, changes in pattern location and the perception of color contrast, and demonstrate how our approach resolves these challenges.

## RESULTS

### Bumble Bee (*Bombus*)

To test the validity of our approach when standardized color templates are used, we applied our analytical approach to a set of bumblebee color templates developed by Williams (2008). Bumble bees exhibit exceptional color diversity largely as a result of mimicry (Williams 2007, Ezray et al. 2019). Some bumble bee species are socially parasitic (~31/260 spp.) on other bumble bee species, as they infiltrate the nests of other species and force host workers to rear their offspring for them. These social parasites thus have established a close association with certain host species, with variable host breadth depending on the social parasite. This association is hypothesized to drive parasites to mimic the coloration of their hosts (Lhomme and Hines 2018). Williams (2008) used color templates to develop criteria for classifying color pattern groups which were then used to test if bumblebees of the obligate, social parasitic subgenus, *Psithyrus*, resemble their host bumblebees. The criteria established included discrete categorization of tail color, pale band color, and pale band position (Williams 2008). Through the application of a randomization test on these characters, Williams (2008) concluded that color patterns of parasite-host pairs in Europe were significantly more similar than would be expected by chance, while parasite-host color pattern association was not significant in North American pairs (Williams 2008).

We applied our approach directly to the colorized templates of Williams (2008) to quantify the perceptual distance between all host-parasite pairs in Europe and North America. We then compared the distribution of distances between host-parasite pairs to the distribution of distances between non-host-parasite pairs following the same analytical procedure as Williams (2008) (Fig 1). The resulting analysis supports the original conclusions of Williams (2008) that in Europe, bumblebees of the genus *Psithyrus* more closely resemble their hosts than is expected by chance and that this pattern does not exist in North America.

**Fig. 1:**
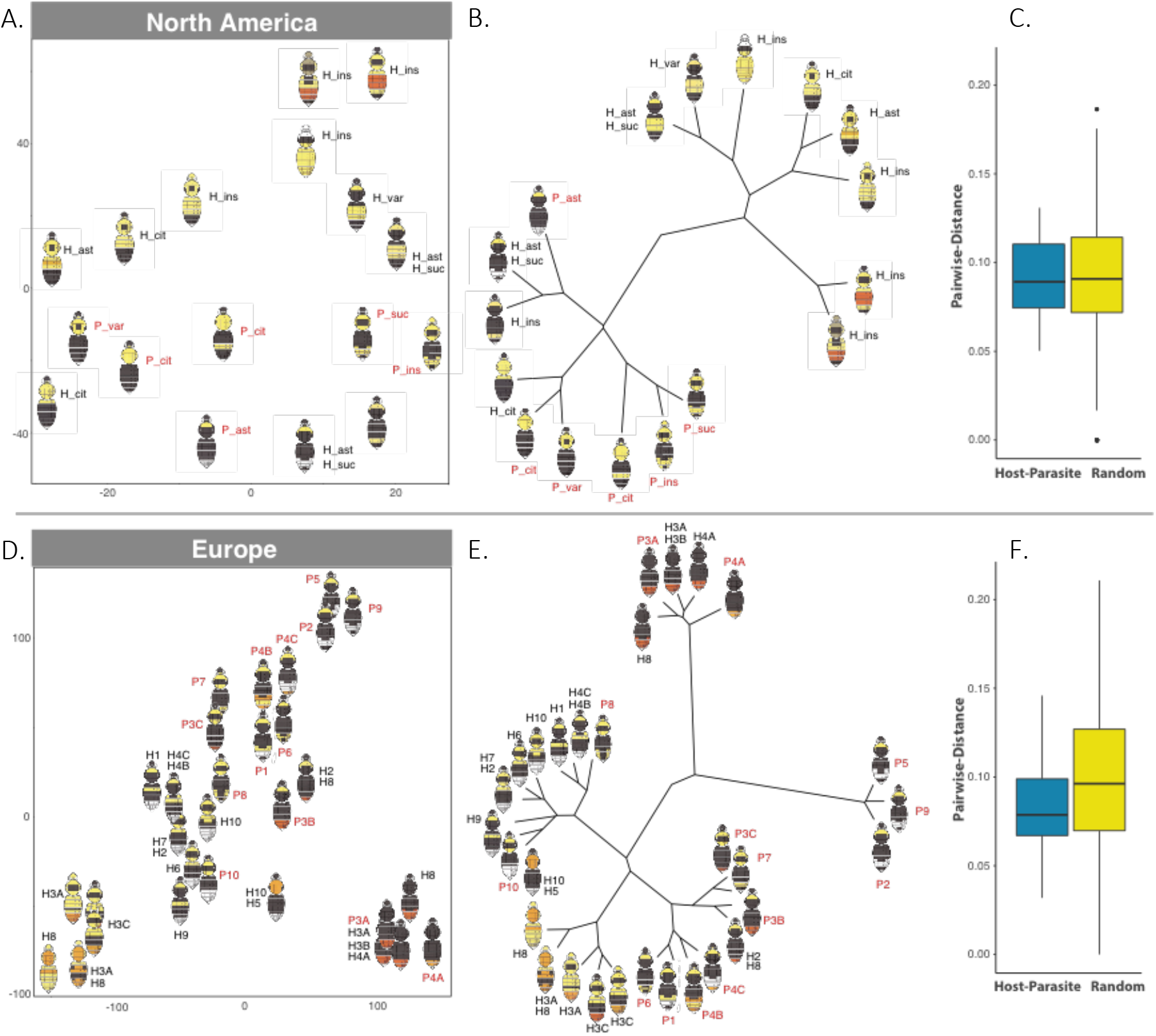
Analysis of bumble bee templates from Williams (2008). Unsupervised analysis of bumble bee host-parasite pairs in North America (**a-c**) and Europe (**d-f**). **a,c** t-SNE plots of perceptual distance with color templates displayed at each embedding coordinate. Host species abbreviations are labeled in black. Parasite species abbreviations are labeled in red. For Europe, host-parasite combinations are recognized by the same number, preceded by “H” for host or “P” for parasite. If a species is a host of multiple species it has multiple identifiers. **b,e** Hierarchical clustering of the perceptual distance of color templates. **c,f** Box plot summarizing LPIPS between host-parasite pairs and random pairs.

We were able to explore continuous aspects of color pattern variation in more detail than Williams (2008) by applying t-distributed stochastic neighbor embedding (t-SNE) to the perceptual distance matrix to visualize its structure (Fig 1a and d). To investigate the degree of support that should be given to discrete groups, we also applied our t-SNE based hierarchical clustering approach to the pairwise perceptual distance matrices (Fig 1b and e). Our results suggest that bumblebee color patterns exhibit clinal variation in multiple dimensions (Fig 1b and e). In Europe, for example, two important axes of variation are observable in the occurrence, position, and size of both a white tail and a yellow stripe in the middle of the body. These two characters were important elements of Williams’ classification criteria (Williams 2008). The t-SNE plot organizes bumble bees that exhibit variation on this color pattern on a cline. Due to the stochastic nature of t-SNE, the position and orientation of this cline varied, but was observable in multiple iterations of the approach (see supplement Fig S1 for 10 sequential random seeds) demonstrating that the approach is able to capture the continuous nature of this variation. Similar to Williams (2008), our results show perceptual clustering of some closely associated host-parasite pairs, which is likely driven by regional Müllerian mimicry groups (Ezray et al. 2019, Williams, 2007). However, individuals cluster by phylogeny as well (Hines and Cameron 2010), as the monophyletic social parasites often cluster together rather than with their hosts in both continents.

### Heliconius

Butterflies present great templates for examining color, as their wings are essentially twodimensional structures and are pattern rich. The images of 28 mimetic butterflies of the Genus *Heliconius* derived from Eltringham’s (1916) color plate XII have been analyzed previously using a variety of methodological approaches (Endler 2012, Maia et al. 2019), and thus make a good system for methodological comparison. Eltringham’s (1916) butterflies are organized into 14 model-mimic pairs, providing an ideal validation set for color pattern analysis as a good perceptual metric should return a lower distance between model-mimic pairs than random pairs. Endler (2012) and Maia et al. (2019) demonstrated the validity of their analytical framework by testing this hypothesis. Their approach begins by applying K-means clustering to the original templates, to reduce the diversity of colors into discrete color categories. Then, a set of color pattern parameters were inferred from this lower dimensional representation which accounts for the relative area of color patches as well as the relative frequency of color transitions and adjacency. Endler (2012) and Maia et al. (2019) both demonstrated that a distance metric derived from their color pattern parameters was indeed lower in model-mimic pairs than non-model-mimic pairs. Endler (2012) also created an unweighted pair group method with an arithmetic mean clustering tree from the color pattern parameters demonstrating that model-mimic pairs clustered together in most cases.

We applied our approach to images derived from Eltringham’s (1916) original plates with minimum processing to remove text and ensure uniform background. Our approach grouped 12 of the 14 described pairs as sibling end nodes with those pairs occurring in > 95% of bootstrap replicates (Fig 2a). Similar to the results of Endler (2012), our approach did not support pair 25-26 as clear perceptual pairs, relative to the other butterflies presented here. Broadly, our results match those of Endler (2012) and Maia et al. (2019). Our method gives better resolution to the variation in model-mimic pairs 1, 2, 7, 10 and 14. However, our method did not support the grouping of model-mimic pair 7 that Endler found in his (2012) analysis. We attribute this to the K-means clustering of colors used by both Endler (2012) and Maia et al. (2019) during the preprocessing of the original images which reduced the number of colors present in the original plates to 4 uniform colors for all images. To investigate the contribution of this preprocessing step to the few discrepancies we observed between our results and those of Endler (2012) and Maia et al. (2019), we re-ran our analysis on the cartoonized plates used by Maia et al. (2019). Our approach grouped all 14 pairs described by Eltringham’s (1916) on Maia et al.’s (2019) cartoonized plates with some minor variation in bootstrap support (Fig 2b).

**Fig. 2:**
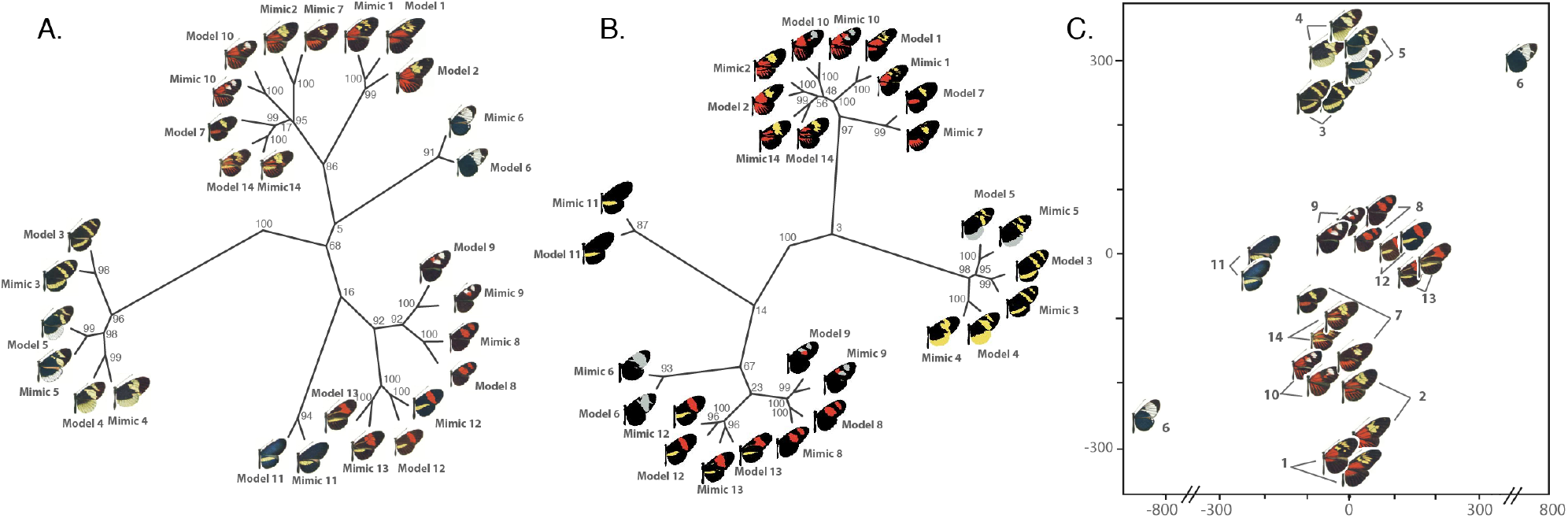
Unsupervised support of Eltringham’s (1916) mimicry pairs. **a-b**, Hierarchical clustering analysis of Heliconius butterfly model and mimic pairs using (**a**) images of Eltringham’s original plates (**b**) the cartoonized templates used by Maia et al. (2019) and (c) t-SNE based plots of perceptual distance of images from Eltringham’s original plates.

### Velvet ant

Templates can take considerable time to build from scratch and involve some subjectivity and data removal in their simplification. Automated approaches such as those used for butterflies work on largely two-dimensional surfaces, but are more difficult when the organism is threedimensional. To test how the approach works on images of pinned, three-dimensional, specimens, we utilized images of North American velvet ants originally published by Wilson et al. (2015). The stinging velvet ants exhibit bright aposematic color patterns and diversity also likely driven by Müllerian mimicry (Wilson et al. 2015). Wilson et al. (2015) used a priori assessment of geographic region, prior research, and visual similarity judged by human observers to propose that North American velvet ants cluster into 8 mimicry rings. Wilson et al. (2015) developed 14 morphological color and color pattern characters that were used to score 351 species of North American velvet ants. Additionally, the authors also applied latent class analysis to these features, grouping them into 8 latent classes and applying a supervised dimensionality reduction technique (nonmetric multidimensional scaling, NMDS) to rescale them by their importance in discriminating between these 8 mimicry rings. While many members of the proposed mimicry rings were grouped into the same latent class, many mimicry rings contained members that were grouped into latent classes with members of other mimicry rings (Wilson et al. 2015). This suggests that mimicry rings may not be fully discrete entities, rather members of seperate rings may frequently share aspects of variation.

We utilized images of 5 specimens per mimicry ring, selecting the 5 that most closely matched the central point for each mimicry ring in Wilson et al. (2015)’s NMDS analysis. We tested the hypothesis that perceptual distance between members of the same mimicry ring should be lower than between mimicry rings. Our results support this hypothesis, with patterns clustering by mimicry complex in the t-SNE analysis (Fig 3). Our unsupervised approach, as well as Wilson et al. (2015)’s, did not show perfect discrete cohesion to the proposed mimicry ring groupings (Fig 3b). Our analysis avoided categorical assignments and thus was better able to represent the continuous nature of variation. We found that patterns form a geographic gradient, with adjacent geographic regions showing more similarity to each other in color pattern, supporting the idea that mimicry patterns can fall along a continuum (Ezray et al. 2019) and that some mimics can straddle mimetic groups through imperfect mimicry of each pattern (Wilson et al. 2013).

**Fig. 3:**
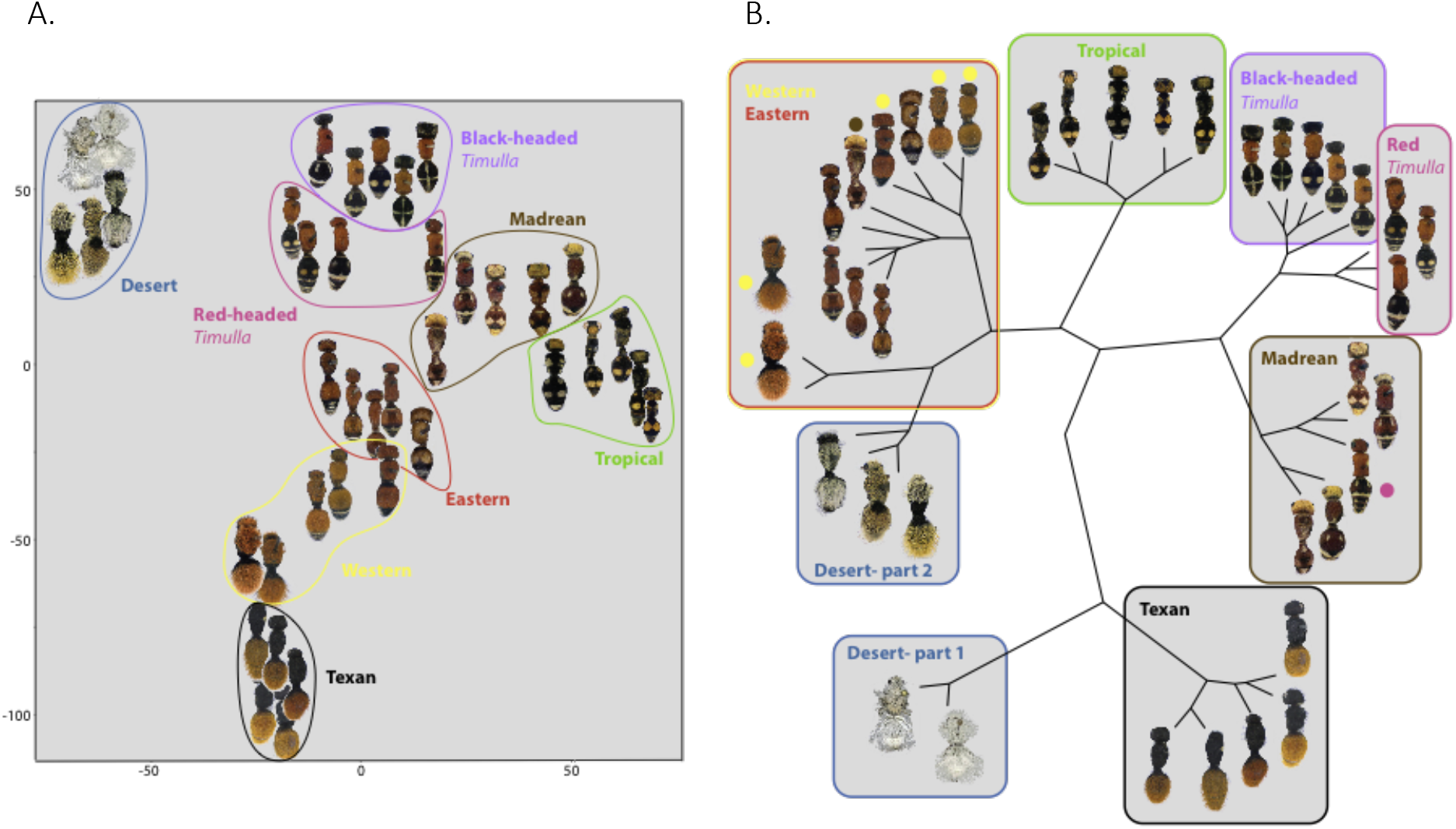
Unsupervised support of previously described mimicry rings of velvet ants. **a,** t-SNE based plots of perceptual distance of velvet ant mimicry rings with the mimicry ring assigned by Wilson et al. (2015) outlined and labeled. **b**, Hierarchical clustering analysis of the same velvet ant images marked and labeled with assigned mimicry rings from Wilson et al. (2015). Individual images are marked with a colored dot if they belong to a different mimicry ring than their circumscribed group.

### Guppy

To further assess the application of our approach on standardized three-dimensional images, rather than templates, and consider their precision for other applications besides mimicry, we utilized two datasets from a genetic study of pigment pattern formation in guppies (*Poecilia reticulata*) (Kottler et al. 2013). The first dataset contains 128 images of F1 males from 4 experimental crosses of wild males from 3 populations mated to a common female population. The resulting dataset consists of a set of sibling males from Cumana and Guanapo and 2 families of sibling males from Quare. This experiment was designed to examine Y-linked color pattern elements of each population. The second dataset consists of images of the same fish taken every 3 days during development, from birth to sexual maturity. These images were taken for 8 separate fish, 6 males and 2 females.

In their original study, Kottler et al. (2013) did not develop a color pattern metric from these two image datasets. Kottler et al. ‘s (2013) experimental design, dataset structure, and conclusions, however, suggest several testable hypotheses. First, we tested the hypothesis that F1 males from different paternal populations should have a larger perceptual distance than males from the same population. Additionally, because the dataset contained 2 families from the Quare population, we also tested the hypothesis that males from different parents should have a greater perceptual distance than sibling males. We then utilized the second dataset to test if perceptual distance tracks developmental time.

Our approach differentiated F1 males from the 3 original populations with strong bootstrap support (Fig 4a). Males from the 2 Quare families were statistically indistinguishable in our analysis (p=0.5422, t-test on the pairwise distances of individuals). t-SNE plots and hierarchical clustering suggest, however, that there are perceptual subgroups within the Quare population. In our analysis, these subgroups contained both siblings and non-siblings suggesting that discrete heritable traits might be responsible for the formation of these subgroups.

**Fig. 4:**
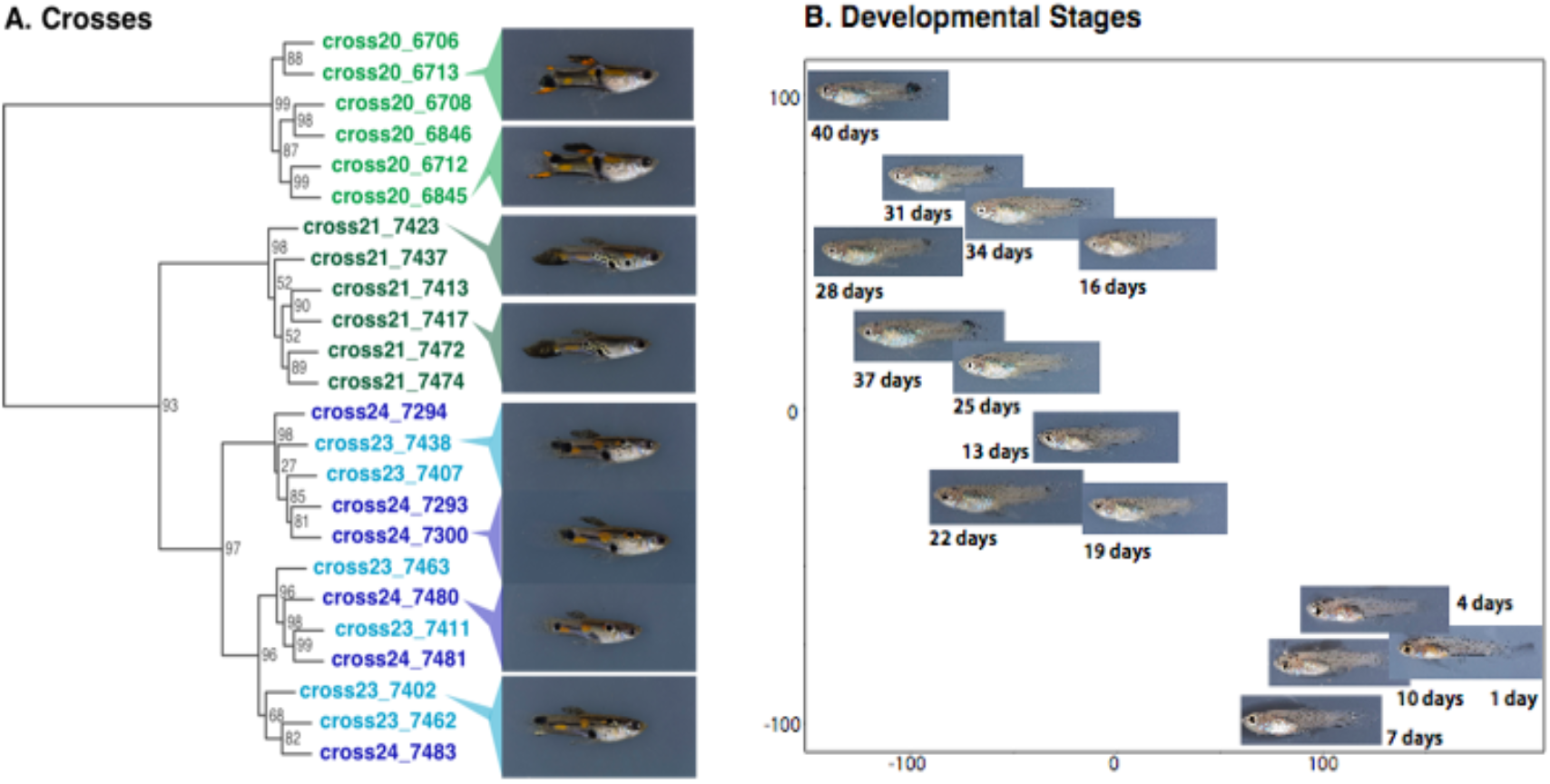
Unsupervised clustering of populations and organizing images along a developmental cline. **a,** Hierarchical clustering analysis of F1 males from 3 different populations. Siblings in cross 20 share a common father from the Cumana population, cross 21 siblings share a common father from the Guanapo population and siblings from cross 23 and 24 come from 2 different fathers from the Quare population. **b**, t-SNE based plots of perceptual distance of images of the same guppy taken every 3 days after birth.

To investigate the relationship between perceptual distance and development, we plotted the relationship between the perceptual distance between images of the same fish and the number of days between the photos (Fig S1). This plot showed a highly significant relationship between perceptual distance and time between images (p=2e-16, generalized linear model), demonstrating that the perceptual metric correlates well to the developmental changes occurring in the guppies in their first 40 days. Similarly, this relationship can be observed in the t-SNE plots of individual fish across all developmental time points which frequently organize themselves on a cline, with similar time points near one another. Hierarchical clustering, however, did not support groupings likely due to the fact that the variation was continuous rather than discrete in nature. (Fig. S1 and see discussion).

### Exploring Pattern Positions with Generated Seahorses Images

Organisms frequently exhibit color patterns that follow a common motif among individuals, but display variability in position. Examples include bands, spots and mosaic color patterns among many others. This type of variation poses a particular challenge to most color pattern analytical techniques because the positional changes in color pattern elements can result in large pixel-wise difference between images while remaining nearly imperceptible. To examine how our approach handles differences of this type, we generated an artificial dataset of banded seahorses. These cartoonized seahorses were designed to show variation in a banding pattern with both in-phase and out-of-phase (bands shifted one band from in-phase) variants. We calculated the pairwise distance between images using pixel-wise L2 distance, a higher level distance metric that incorporates differences in luminance, using a contrast and structure based technique called SSIM (Wang et al. 2004), and the LPIPS perceptual approach. We then followed our proposed t-SNE and hierarchical clustering approach for all three distance matrices.

t-SNE based hierarchical clustering of L2 distance resulted in groups with high bootstrap support separating those with in-phase stripes from those with out-of-phase stripes, and secondarily dividing base color and pattern (Fig. 5). The same analysis utilizing SSIM distance resulted in high bootstrap support groupings by base color, a result that may seem more rational than the results of L2 distance (Fig. S2). However, within base color clusters, out-of-phase striping patterns were grouped separately from their in-phase counterparts (Fig. S2). In contrast, t-SNE based hierarchical clustering using LPIPS resulted in high bootstrap support groupings by base color (Fig 5). Within base color clusters, non-striped seahorses were grouped outside of both their in-phase and out-of-phase striped counterparts similar to how it would be perceived by eye (Fig 5).

**Fig. 5:**
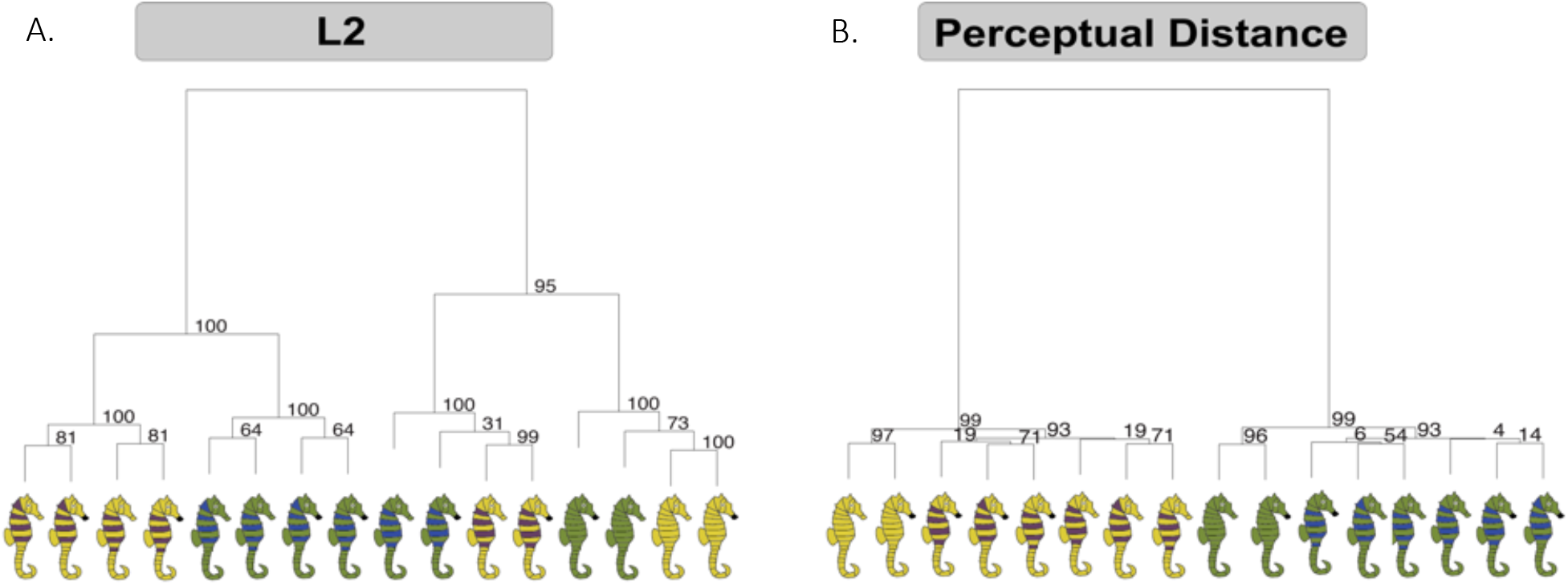
Exploring Pattern Positions with Generated Seahorses Images. **a,** Hierarchical clustering analysis of generated seahorse images based on L2 distance. **b**, Hierarchical clustering analysis of the same generated seahorse images based on LPIPS.

### Exploring contrastive elements with Generated Fly Images

Organisms also frequently display inter-individual or inter-specific variation in color pattern elements that are contrastive against their base body color. Examples include the yellow, orange, and red stripes on the abdomen of bumble bees previously analyzed here (Williams 2007). To examine how changes in saturation and hue relate to distance metrics, we generated two similar datasets of fly templates. Both datasets utilized a black base colored fly template, however, they differed in that one featured a contrastive band on the abdomen and the other featured a contrastive spot on the thorax. These contrastive elements were then varied across regular intervals of the RGB spectrum and analyzed separately.

t-SNE based hierarchical clustering of L2 and SSIM distance organize individuals such that images of similar stripe saturation were near one another in the tree. t-SNE based hierarchical clustering using LPIPS instead recognizes these colors to occur on opposing axes, thus better distinguishing among color types first and then saturation level (Fig 6).

**Fig. 6:**
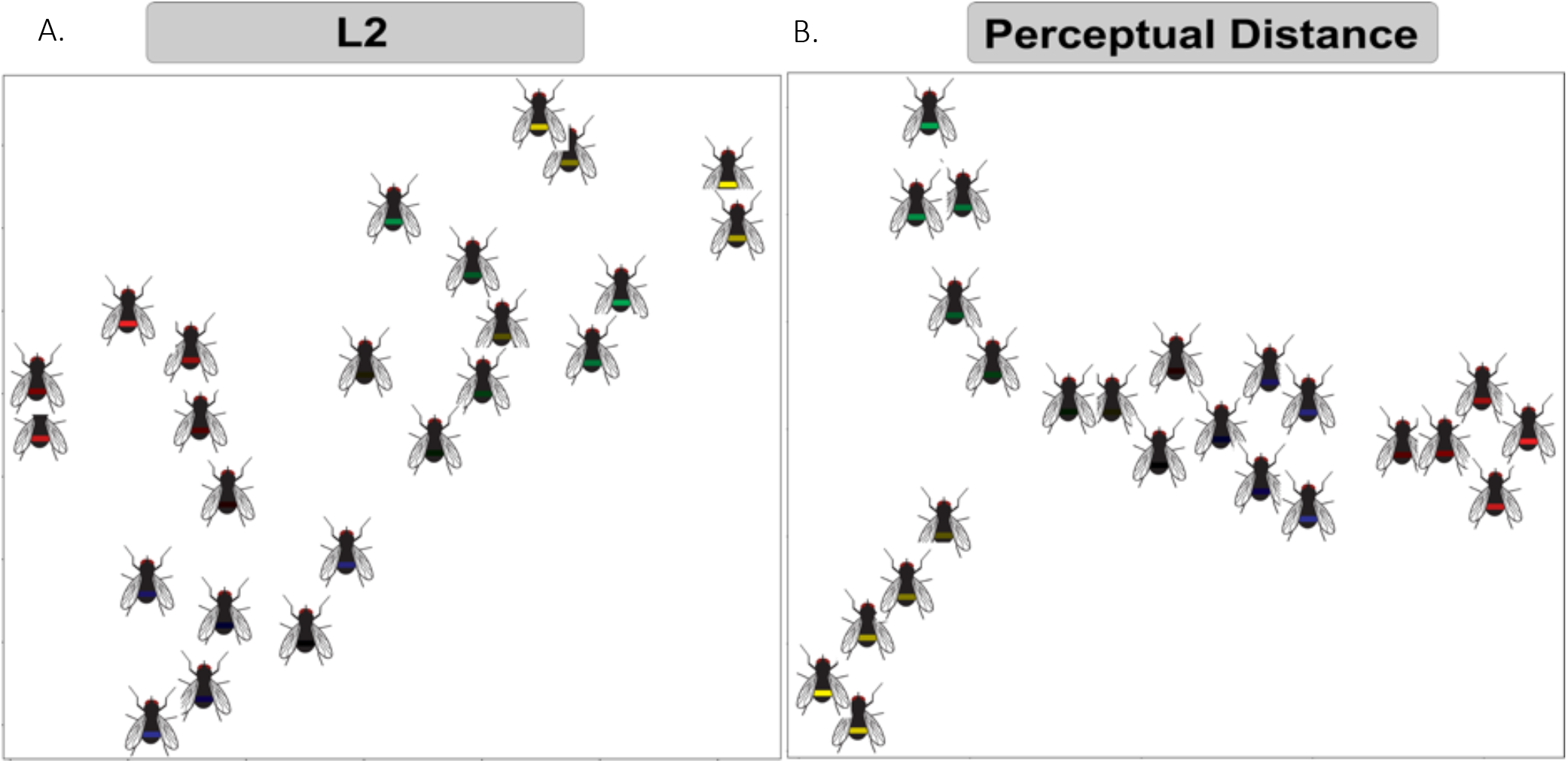
Exploring contrastive elements with Generated Fly with Stripes Images. **a,** t-SNE based plots of generated fly templates based on L2 distance. **b**, t-SNE based plots of generated fly templates based on LPIPS.

## DISCUSSION

Here, we present a pipeline for quantitative analysis of organismal color pattern that utilizes the features of a pre-trained, deep CNN to generate perceptual distance. We validated our approach by applying it to previously published data. These datasets varied in their taxonomic focus, as well as the nature and amount of variation they contained. Additionally, these datasets also ranged from cartoonized templates, to drawings, to images. Despite the diversity of application, our approach was able to reproduce many of the previously reported findings from these datasets. Importantly, these results were achieved without task specific training, the creation of classification criteria, or trait matrices.

We believe these aspects of our approach offer significant advantages in a variety of contexts, however, applied in the incorrect context these strengths become weaknesses. Here, we examine the contexts in which this approach is best applied, as well as, the context in which an alternative supervised, semi-supervised, or non-machine learning method might be a better choice. We also discuss experimental design considerations for any future work that wishes to utilize this analytical approach.

### Classification Criteria

A common approach for examining color pattern variation is to develop discrete color pattern traits and then score a set of samples or species based on the presence and absence of these traits. Both the bumble bee (Williams 2008) and velvet ant (Wilson et al. 2015) datasets were previously analyzed under this general framework. In both cases, criteria were developed based on each set of researchers observations of important axes of variation in their respective system. It may be useful or necessary to participate in the criteria or feature selection process for several reasons, including having specific interest in only some aspects of color pattern or interest in regressing some other variable against the trait matrix. This approach yielded discriminative criteria for groups in both studies that related to other aspects of the organisms biology.

It may in some cases be important, however, that clusters be defined through unsupervised methods. The existence of mimicry rings, for example, is often proposed by demonstrating statistical support for visually similar clusters of organisms. Unsupervised clustering methods can be applied to the classification criteria or trait matrices after they are scored, as Wilson et al. (2015) did in their analysis. However, this approach is not fully unsupervised because the researcher provides the perceptually salient criteria that form the basis for the inferred clusters. In these cases, researchers might find value in our approach as a fully unsupervised method for independently supporting the existence of visually similar groups.

### Advanced computational techniques

Numerous computational techniques have been designed to account for many of the aspects of quantifying color pattern similarity. Many of these techniques, which include boundary strength analyses (Endler et al. 2017), adjacency analyses (Endler 2012), and spectral clustering implemented in advanced frameworks such as PAVO 2 (Maia et al. 2019) and Patternize (Van Bellegham et al. 2018) have analytical elements that are designed to address many aspects of color pattern quantification addressed by the method we present here. The convolutional nature of our analytical approach, however, allows for a more complete realization of these original ideas. In these programs, the relative frequency of transition between colors (adjacency analysis), as well as the position and strength of contrast boundaries (boundary strength analysis) among other facets of color pattern variation, are developed into a feature matrix. These aspects of color pattern are accounted for in the method we present here at every perceptual scale. Additionally, a representation of color pattern similarity is created through the original training of the network on the classification task, as well as, in the human judgment based re-weighting of features (Zhang et al. 2018). This removes any need to reduce the colors to factors through K-means clustering. Finally, CNNs were designed to be positionally/translationally invariant, meaning that the representation of objects is robust to minor movement in the frame, thus negating the need for precise image alignment. We, however, suggest that researchers still align and scale their images to reduce unnecessary sources of variation.

Here, we demonstrate that deep feature based distance is able to capture many of the same perceptual elements as those addressed by boundary strength and adjacency based analyses. We validated this finding by comparing the results of our analysis of Eltringham’s (1916) *Heliconius* mimicry pairs to previously published results of the same dataset. These results were achieved by applying our methodology directly to images derived from the original color drawings, with no changes to color, cartoonization, or algorithmic alignment of the color plates. Our method did not support grouping pairs 2 and 7 in the hierarchical clustering analysis when images derived directly from the plates were used, although these were in close proximity to each other in the t-SNE plot. Close examination of the original color plates revealed that the zone of overlap between the forewing and hindwing, contains a brown patch in only some of the butterflies. In Endler (2012) and Maia et al. (2019)’s analyses this region was homogenized by the application of K-means to reduce the continuous colors of the original plates to a smaller set of discrete colors which removed this feature from being considered as an axis of variation. This appears to explain why these butterflies are clustered together in our analysis of the original plates, but they were not clustered when we analyzed the cartoonized plates used by Endler (2012) and Maia et al. (2019) (Fig. 2). Researchers may wish to use image pre-processing software to cartoonize images prior to using the perceptual metric described here when elements of variation along these minor color and patch variation are not of interest. When these elements of variation are of analytical interest, the present methodology is uniquely capable of integrating and weighting these elements along with all other elements of perceptual differences.

### Utility of the Method for Photographs

We also investigate the utility of deep feature based perceptual metrics applied directly to photographs. Compared to template based and cartoonized datasets, photographic datasets contain additional aspects of variation which include slight differences in pose of the subject, minor differences in lighting, greater heterogeneity of color among many others aspects of variation. While researchers should strive to minimize these sources of variation, wherever possible, we demonstrate that deep feature based perceptual distance is robust to many of these sources of variation.

In our analysis of Kottler et al.’s (2013) guppies, we did remove images from our analysis that contained large artifacts such as shadows and reflections. However, there was minor heterogeneity in the position of the fish, as well as lighting of both the fish and the blue background. These sources of variation likely contributed to the high variance observed in the relationship between perceptual distance and the number of days between images (Fig. S1). Yet, despite this heterogeneity, deep feature-based perceptual distance still recovered the biologically meaningful patterns of population differentiation (Fig. 4), as well as, clinal variation in perceptual distance along the developmental timeline (Fig.4 and S1).

Similarly, in our analysis of Wilson et al.’s (2015) velvet ants, the original images contained significant differences in leg positioning as well as the presence, color, and positioning of the pin holding the mounted specimens. To reduce this source of heterogeneity, we removed the legs from the images, however, we were unable to address differences in the presence and placement of pins, which likely contributed to our final results. Despite these issues, however, our analysis grouped velvet ants into similar mimicry groups to Wilson et al. (2015). We anticipate that if future studies are designed with the intention of using this technique, these extraneous sources of variation can be minimized during data collection.

### t-SNE and Hierarchical Clustering Visualizations

In addition to developing a pipeline to produce a perceptual distance matrix for a given dataset based on LPIPS, we also developed several visualization and analytical methods. These methods are designed to support the analysis of a wide variety of relationships between color pattern variants in a dataset, including both discrete and continuous variation. The unsupervised technique t-SNE provides an important aspect of this workflow. The application of this technique converts the high-dimensional, non-euclidean, perceptual distance matrix to a set of coordinates for each image. While reducing the dimensionality of the dataset, t-SNE attempts to preserve the structure that exists between groups in higher dimensions. Elements of variation that are shared between some images, but not others, are therefore weighted higher. This results in an important property which we feel is consistent with a perceptual concept. When variation occurs over one or more perceptual axes those attributes should be organized on a manifold, however, when differences are sufficiently large with no intermediate forms those variants should be placed arbitrarily far away from one another. Therefore, additional differences do not result in additional distance in this context. Thus, this approach preserves both local continuous variation (i.e. guppy development) as well as large scale discrete clustering (guppy population differentiation).

It can, however, be difficult to distinguish when patterns in t-SNE space are due to actual patterns in the underlying perceptual distance matrix or if they are due to the stochastic element of t-SNE (Wattenberg et al. 2016) (e.g., flattening multidimensional data placed mimicry pairs in opposite corners in Fig. 2C even though they cluster closely in the tree). For this reason, we developed a bootstrapped UPGMA tree based method which iterates t-SNE and then performs hierarchical clustering on the consistency of observed patterns in t-SNE over independent t-SNE analyses of the same data. The frequency in which groups appear consistently across independent t-SNEs is interpreted from the resulting trees which is useful when considering the degree of support for discrete groups. This approach, however, is less effective in assessing support of similarity in the presence of continuous variation. Similarly, phylogenetic clustering fails to recognize that some patterns are intermediate between others. For images arranged along a cline (Fig. 4), this pulled taxa apart even when they occur in close proximity in the t-SNE plots and generated low bootstrap values. For this reason, when continuous variation is a significant feature of a dataset, we suggest visualizing the dataset by selecting a representative t-SNE plot rather than using the hierarchical clustering based tree.

### Conclusion

Numerous other approaches have been developed to quantify and analyze organismal color pattern. These approaches include expert-derived trait matrices, metrics designed to capture individual salient aspects of perception, and sophisticated applications that combine many such metrics like the workflows supported by QPCA (van den Berg et al. 2019), PAVO 2 (Maia et al. 2019) and Patternize (Van Bellegham et al. 2018). Many of these applications also can incorporate a particular receiver’s visual acuity and spectral sensitivity into their analysis. These approaches are well suited for studies where (1) the aspects of variation that are of interest are already known, (2) when only one or several salient aspects of perception need to be considered, (3) and/or when the perspective of a particular receiver is the only one of interest.

Current methods, however, are not well suited to accommodate analysis when (1) the important aspects of variation are not known or not easily scored because they are of a continuous nature, (2) all salient aspects of perception need to be considered simultaneously, (3) and/or the relevant receiver is not known or the color pattern is relevant to a diverse set of receivers. These circumstances apply to a broad array of applications in biology and they are most appropriately addressed by an unsupervised analysis based on a general model of perception. The proposed unsupervised approach which utilizes the newly developed perceptual distance metric, LPIPS, was not developed to approximate any salient aspect of perception. Instead, it is a result of training a system to first solve the higher-level task of general classification. A perceptual metric was then developed from the internal representations of this general classification model (Doersch et al. 2015, Zhang et al. 2018). Perception in LPIPS arises from the difficulty of classification or confusion of two groups due to the sharing of highdimensional visual features. Zhang et al. (2018) suggested that perceptual judgment is not a specialized task, rather, it is an emergent property of systems that have been designed or evolved to solve broad semantic tasks. If true, we find the use of a perceptual metric developed in this way particularly compelling in the context of biological investigations. The broad utility of the Perceptual Loss metrics and LPIPS has resulted in rapid adoption and use across an array of computer vision tasks. Similarly, because of its effectiveness in the context of measuring organismal color patterns, we anticipate that variations of the technique will rapidly be applied to a diverse set of biological investigations.

## Methods

### Datasets and image processing

Image datasets used to generate figures are available in our github repository: https://github.com/DrewWham/Perceptual_tSNE. Bumble bee templates featured in Fig. 1 were extracted from Williams (2008) and utilized in analyses here without editing apart from scaling to 256×256 square images. Heliconius images featured in Fig. 2 were derived from Eltringham’s (1916) color plate XII. We centered and scaled each butterfly into its own 256×256 square image and removed any text to ensure background uniformity. Velvet ant images featured in Fig. 3 were extracted from Wilson et al. (2015) to represent the phenotype of each mimicry complex. As these original images displayed differences in leg positioning, the legs of each specimen were removed to reduce heterogeneity not relevant to color pattern. These cropped images were scaled to 256×256 square images prior to analysis. Guppy images utilized in this study were derived from Kottler et al. (2013). Images that contained large artifacts such as shadows and reflections were removed, the images that were analyzed were rescaled to 256×256 square images prior to analysis.

### Generated Seahorse and Fly datasets

To test the robustness of our approach to changes in position of color pattern elements, we designed the Seahorse dataset with 5 key elements of variation; (1) two base color groups each containing contrastive color variation in alternating segments to create a banding pattern, (2) the green (rgb: 119,143,59) and yellow (rgb: 220,203,50) base colors were selected to be more similar to each other numerically than to the blue (rgb: 52,78,161) and purple (rgb: 103,65,99) contrastive colors, (3) start the banding pattern at 3 separate segments such that 2 patterns are in-phase and 1 is out-of-phase, and (4) variation in a minor visual element (dark colored first segment). Variations across these elements resulted in a 16 image dataset.

To test the importance of contrastive elements as well as the effect of variation in their contrast from the background color, we designed two similar fly datasets. These datasets were identical in design, however, in one the contrastive element was a dot in the middle of the flies thorax and in the other, the contrastive element was a stripe across the flies abdomen. Each of these datasets contained variation in the contrastive elements with four base colors: red (rgb: X,0,0), green (rgb: 0,X,0), blue (rgb: 0,0,X) and yellow (rgb: X,X,0). The non-zero RGB channel values were varied across 6 levels (255, 204, 153, 102, 51, 1) for each base color. Variations across these elements resulted in 2 different 24 image datasets.

We analyzed each of these datasets by calculating a pairwise distance matrix using L2 distance, SSIM distance, and LPIPS. We then applied the analytical techniques described in the following section to each of these pairwise distance matrices.

### CNN implementation and post-model analyses

All input images were resized using the skimage package in python (van der Walt et al. 2014). We then utilized Zhang et al.’s (2018) implementation of LPIPS to extract the activations of the pre-trained AlexNet CNN (Krizhevsky et al. 2012) and applied a linear rescaling of these activations. This yielded a pairwise perceptual distance matrix for a given dataset of images. We then derived a separate distance matrix from the perceptual distance matrix using t-distributed stochastic neighbor embedding (t-SNE) (van der Maaten 2008, van der Maaten and Hinton 2014) implemented in the R tsne package (Krijthe et al. 2018). This method stochasticly generates a 2-dimensional embedding for every image in the perceptual distance matrix by minimizing the Kullback-Leibler divergence between the LPIPS distance matrix and t-SNE distance matrix. To evaluate the significance of hierarchical groups observed in the t-SNE analysis, we then derived 10,000 t-SNE distance matrices from the LPIPS based distance matrix. Furthermore, we applied a hierarchical clustering approach with bootstrapping implemented in the R pvclust package to each of these t-SNE distance matrices (Suzuki and Shimodaira 2006). We then graphed the most common tree structure, reporting for each node the frequency with which all terminal nodes were members of that node across the 10,000 trees. This method is therefore biased towards lower bootstrap values as you move away from the terminal branches, just as other bootstrap methods are, because the more terminal branches have fewer members. The approximately unbiased approach implemented in pvclust (Shimodaira 2004), however, cannot be applied here. These metrics are derived by resampling features and then re-generating a distance matrix from the resampled data to see how sensitive clusters are to perturbations in the number of features used to generate the tree. Our data differs, however, from the character based genetic data that this method was designed to analyze. The approximately unbiased bootstrapping approach assumes that each feature should have equal weight in determining clusters and that bootstrap and p-values should summarize the degree to which features agree on cluster relationships as well as the degree to which the number of features measured is enough to support conclusions. Here, such an approach to generate bootstraps is not meaningful because the features are scaled by their perceptual importance. The elements of uncertainty that are summarized by the bootstrap values here are the stochastic nature of the dimensionality reduction technique and the tree building algorithm. The final tree, bootstrap values, and images are then plotted using the ggtree package (Guangchuang et al. 2017).

---

**Supplemental Figure 1:**
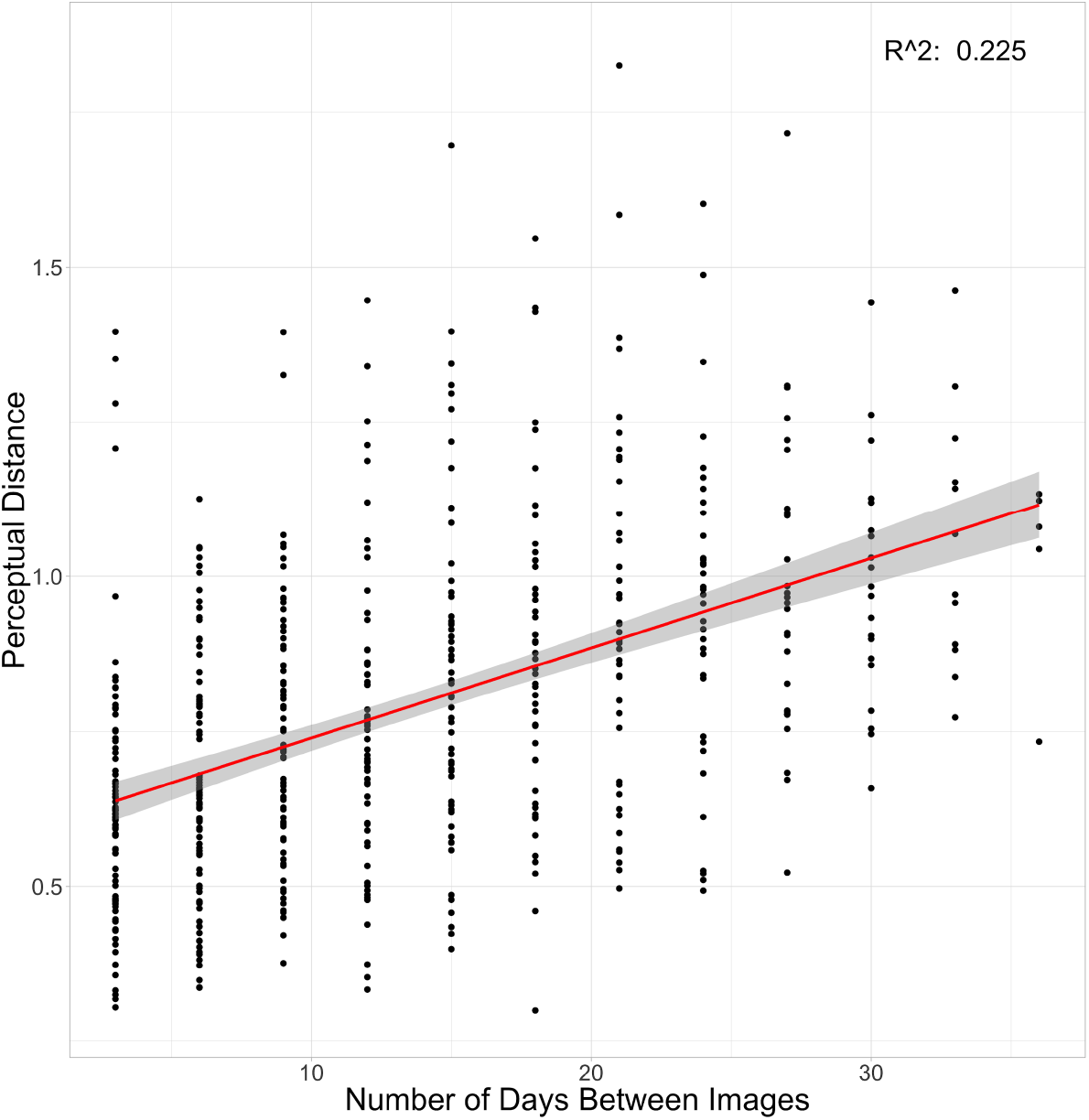
Linear regression between the length of time (days) between when images of guppies were taken and perceptual distance.

**Supplemental Figure 2:**
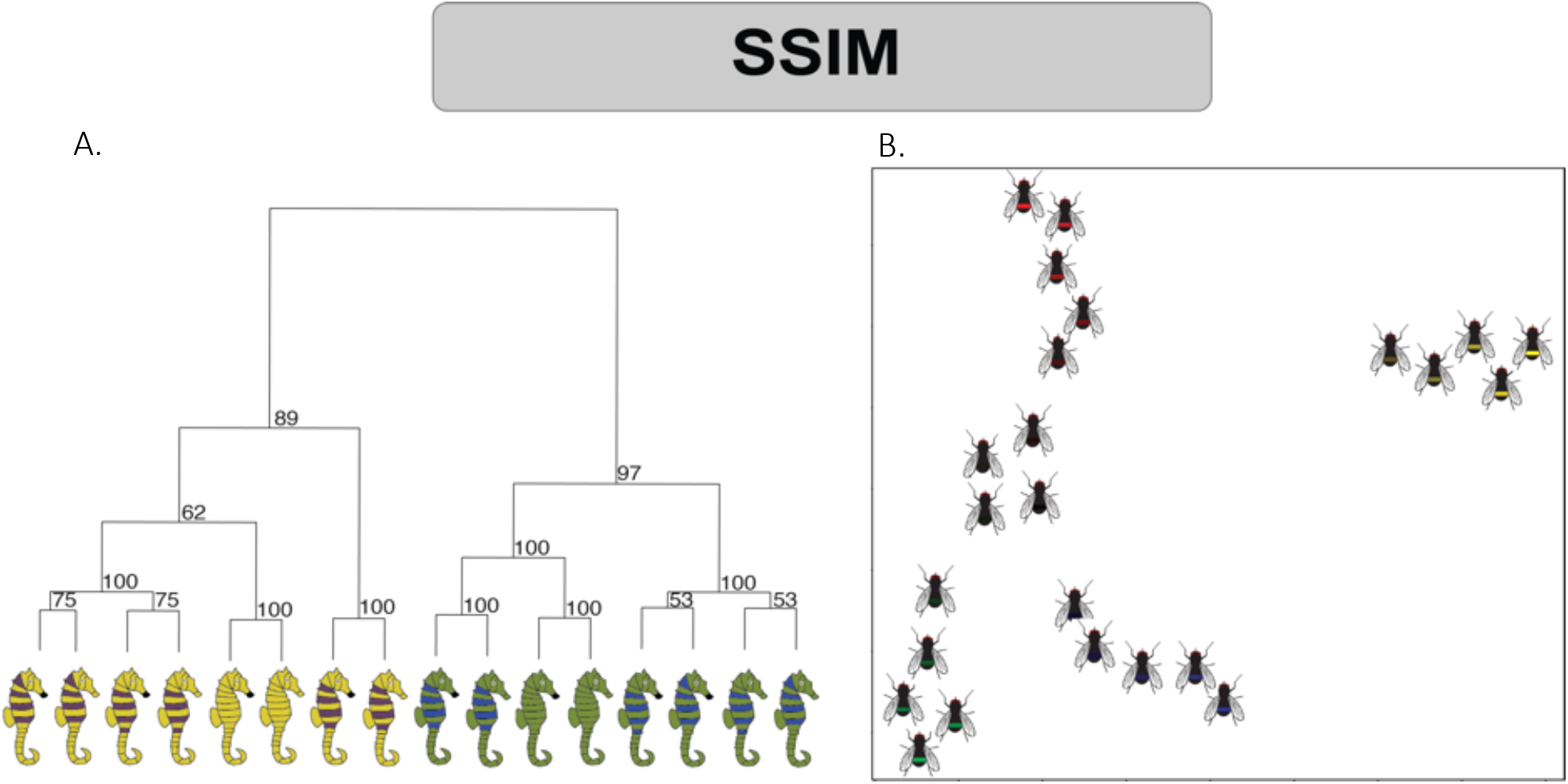
Exploring Pattern Positions with Images of Generated Seahorses and Flies with Stripes Using SSIM. **a,** Hierarchical clustering analysis of generated seahorse images based on SSIM distance. **b**, t-SNE based plots of generated fly templates displaying stripes based on SSIM.

**Supplemental Figure 3:**
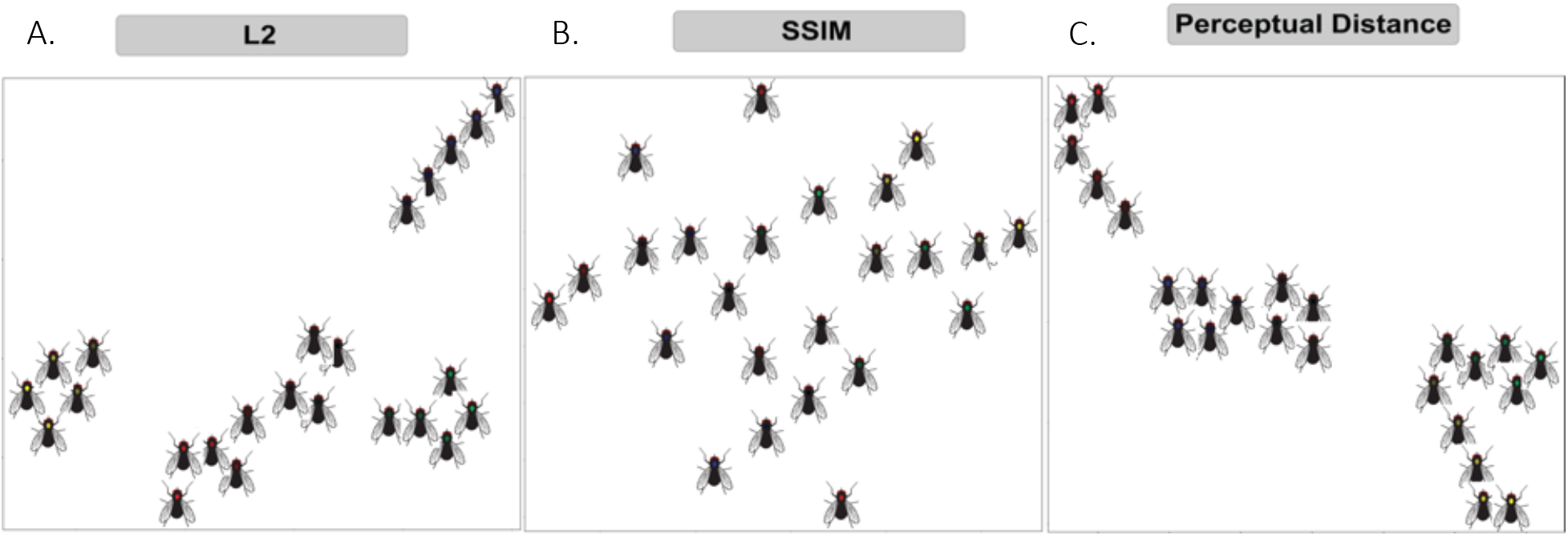
Exploring contrastive elements with Generated Fly with Dots Images. **a,** t-SNE based plots of generated fly templates based on L2 distance. **b**, t-SNE based plots of generated fly templates based on SSIM. **c**, t-SNE based plots of generated fly templates based on LPIPS.

